# Comprehensive evaluation and practical guideline of gating methods for high-dimensional cytometry data: manual gating, unsupervised clustering, and auto-gating

**DOI:** 10.1101/2024.08.12.607667

**Authors:** Peng Liu, Yuchen Pan, Hung-Ching Chang, Yusi Fang, Xiangning Xue, Jian Zou, Jessica M. Toothaker, Oluwabunmi Olaloye, Eduardo Gonzalez Santiago, Black McCourt, Vanessa Mitsialis, Pietro Presicce, Suhas G. Kallapur, Scott B. Snapper, Jia-Jun Liu, George C. Tseng, Liza Konnikova, Silvia Liu

## Abstract

Cytometry is an advanced technique for simultaneously identifying and quantifying many cell surface and intracellular proteins at a single-cell resolution. Analyzing high-dimensional cytometry data involves identifying and quantifying cell populations based on their marker expressions. This study provided a quantitative review and comparison of various ways to phenotype cellular populations within the cytometry data, including manual gating, unsupervised clustering, and supervised auto-gating. Six datasets from diverse species and sample types were included in the study, and manual gating with two hierarchical layers was used as the truth for evaluation. For manual gating, results from five researchers were compared to illustrate the gating consistency among different raters. For unsupervised clustering, 22 tools were quantitatively compared in terms of accuracy with the truth and computing cost. While no method outperformed all others, several tools, including PAC-MAN, CCAST, FlowSOM, flowClust, and DEPECHE, generally demonstrated strong performance. For supervised auto-gating methods, four algorithms were evaluated, where DeepCyTOF and CyTOF Linear Classifier performed the best. We further provided practical recommendations on prioritizing gating methods based on different application scenarios. This study offers comprehensive insights for biologists to understand diverse gating methods and choose the best-suited ones for their applications.

## INTRODUCTION

Cytometry is a powerful single-cell assay that allows for high-dimensional profiling of diverse cell populations in suspension^1–3^. This technique has been widely applied to clinical diagnosis, immunology and cancer research, and the pharmaceutical industry^3–5^. Flow cytometry (FCM) and mass cytometry (or Cytometry by Time-Of-Flight, CyTOF) are two major techniques that employ labeled antibodies to quantify the cell surface and intracellular proteins. FCM labels the markers by fluorescence and measures the emitted fluorescence emitted per cell as they pass individually through a laser beam. The traditional FCM technique can detect around 8-10 markers, while the recent study has developed a 43-color flow cytometry panel^6^. As an advanced technique, CyTOF utilizes antibodies chelated with heavy metal isotopes to identify cell surface and intracellular proteins. The metal isotopes are primarily from the lanthanide series of elements, making them neither biologically derived nor radioactive. Once cell suspensions are stained and introduced into the mass cytometer, they are nebulized into droplets containing individual cells. The droplets are then ionized with argon plasma to release the metal isotopes attached to the antibodies in each droplet. The ions are separated based on mass such that the lower mass biologically derived atoms are removed and those of higher mass enter a time-of-flight chamber to measure their mass-to-charge ratios, allowing for the quantification of relative isotope abundance in each droplet^7–9^. Since mass cytometry has minimal background and high specificity, CyTOF allows for simultaneous combined measurement of up to 40-50 different antibodies, enabling the identification of a large number of cellular populations from an individual sample.

Analysis of cytometry data, although a powerful technique, brings challenges to the community. One of the major obstacles is to properly identify cell populations among thousands to millions of cells based on their high-dimensional markers^10^. In this study, we performed a comprehensive quantitative evaluation of available gating methods for high-dimensional cytometry data. **Figure 1** provides a flow chart demonstrating the cytometry data format, composition of marker tables, and the evaluation pipeline of three gating categories in this study: manual gating, unsupervised clustering, and supervised auto-gating. The first manual 2-D gating is the most traditional method used by biologists to identify cell populations. As illustrated in **Figure 1**, in the “Manual gating” block, pairwise makers are selected based on prior knowledge (known markers of interest) and applied to identify a subset of cells. This subset of cells can be further selected and grouped by other makers to identify cell subpopulations. Eventually, hierarchical layers of cell populations are built according to different markers applied. Manual gating can be performed using FlowJo (BD Life Sciences), Cytobank (Beckman Coulter), or other software with friendly graphical user interfaces. It has advantages in identifying different cell populations of interest straightforwardly and flexibly. However, the gating process is experience-based, time-consuming, and relies on prior knowledge and arbitrary cutoffs to assign cell populations.

**Figure 1:**
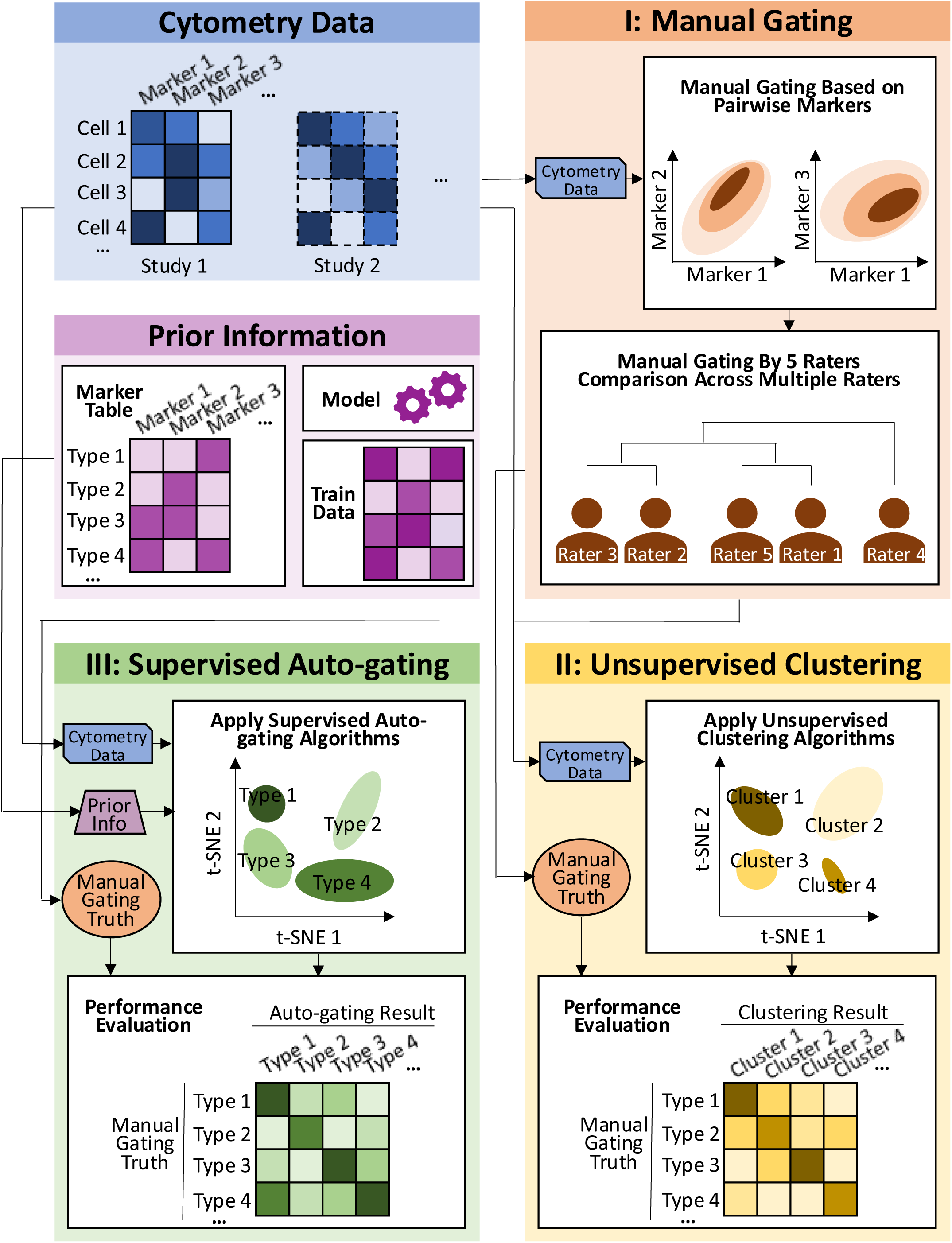
**Workflow of the cytometry data analysis pipeline**. The cell-by-marker intensity table per study was used as the input. For the (I) manual gating method, scatter plots on pairwise markers were drawn to define the cell populations. In this paper, manual gating was performed by 5 raters independently and their gating results were compared. For the (II) unsupervised clustering methods, cytometry data were input into multiple clustering algorithms. Their results were compared to the manual gating (as truth) and evaluated. For the (III) supervised auto-gating methods, both cytometry data and prior knowledge (such as maker table, model, and training data) were used as input. Multiple auto-gating algorithms were applied and evaluated based on the manual gating truth.

In addition to manual gating, computer-aided unbiased algorithms have been developed to identify cell populations in a more automated manner^11^, including the second and third categories: unsupervised clustering and supervised auto-gating (which further includes semi-supervised or supervised gating). For clustering methods, as shown in the “Unsupervised Clustering” block of **Figure 1**, cells are grouped into clusters based on marker intensities without human intervention. Cell type characteristics of the identified clusters are, however, unknown and rely on researchers to further annotate. The supervised auto-gating methods (illustrated in the “Supervised Auto-gating” block of **Figure 1**) not only group cells into clusters based on the marker intensities, but additionally curate the cell populations identified by assigning labels to each cell cluster based on pre-specified cell-type marker tables. Compared to manual gating, computer-aided methods are faster, can simultaneously analyze multiple datasets in a highly efficient and reproducible manner, and do not rely on prior knowledge to cluster the cellular populations. However, these methods sacrifice the flexibility afforded by manual gating.

As shown in Figure 1, three types of gating methods (manual gating, unsupervised clustering, and supervised auto-gating) have been developed with increasing automation and decreasing human intervention. With the advanced popularity and development of cytometry technology, several review and evaluation papers have been published in the last 10 years. Most review literature provides descriptive introductions of clustering and visualization tools and conceptual guidelines to users with little to none-quantitative comparison to support the conclusion^11–20^. To our knowledge, only two papers have performed numerical evaluation. Weber *et al*^21^ compared 18 unsupervised clustering algorithms on 4 CyTOF and 2 FCM datasets. Liu *et al*^22^ evaluated 7 unsupervised clustering methods and 2 semi-supervised methods across 6 datasets to provide guidelines on choosing clustering algorithms for cytometry data. However, a more comprehensive evaluation of tools, especially in manual gating and supervised auto-gating, and extensive panels of evaluation criteria are lacking to conclude a solid guideline for users. In contrast to the limited scope of existing papers, we performed comprehensive investigation and evaluation in all three categories in this paper (see **Supplementary Table 1** for a side-by-side comparison of existing literature and the current paper). Firstly, in manual gating, we collected gating results from five raters in three different labs and evaluated gating consistency across raters. Two hierarchical layers of gating results further served as ground truth to evaluate the gating performance of the other computer-aided algorithms. Secondly, in unsupervised clustering, we attempted 32 unsupervised clustering tools previously reviewed by Liu *et al*^11^ and successfully implemented and compared 22 tools across 6 datasets. Based on the truth from manual gating, we expanded evaluation criteria (adjusted Rand index (ARI) and F-measure) and computing benchmarks to provide an evaluation panel for prioritizing overall tool performance. We also evaluated the tools’ ability to detect rare populations, which is critical in many biological or clinical applications. Finally, in auto-gating, we successfully implemented 4 out of the 6 auto-gating (supervised or semi-supervised) methods^11^ to provide guidelines on the application of automatic cell population identification.

The innovation and merits of this evaluation paper compared to existing papers are highlighted below (**Supplementary Table 1**). (1) <u>Gating methods</u>: Manual gating (5 raters), unsupervised clustering (22 tools) and auto-gating (4 tools) methods were systematically reviewed and compared. To the best of our knowledge, such a comprehensive evaluation has not been performed before, especially since evaluations of manual gating and auto-gating were mostly missing previously. (2) <u>Datasets:</u> Tools were evaluated by both in-house and public datasets, including multiple species (human, mouse, and non-human primates) and cell types (peripheral blood mononuclear cells (PBMC), placental villi, and bone marrow). For in-house data, two hierarchical layers of manual gating were used as ground truth, where the first layer included major populations, and the second layer contained a more detailed identification of sub- populations. Data for two rare populations were also used to check the tool’s ability to detect small clusters of cells. (3) <u>Evaluation benchmarking</u>: Multiple evaluation measurements (F-measure, ARI, Cohen’s kappa index) were employed. Computing time was evaluated on a grid of cell numbers to evaluate the scalability of the tools for ultra-large cell number applications in the future. (4) All programming scripts for tool implementation and comparison were made available on GitHub (https://github.com/hung-ching-chang/GatingMethod_evalutation/). In the research development, we surprisingly found that the implementation of many published tools is not trivial, and many are not achievable after contacting the original authors and attempts by multiple co-authors in this paper. As such, this paper allows future users in this field to easily apply and compare different tools in their datasets. The GitHub resource also provides an evaluation platform when a new gating method is developed in the future.

## METHODS

### CyTOF experimental pipeline for Rhesus macaque (NHP) samples

**<u>Placental samples</u>** were collected from the California Primate Center at the University of California-Davis as described in our published study^23^, and from which CyTOF experiment data was extracted. Briefly, pregnant macaques were injected with either LPS or saline solution and their offspring were delivered via Cesarean section 16hrs following injection and their placental biopsies were collected following delivery. For cryogenic storage, each placental layer sample was stored and processed according to previously published protocol^24^. Fresh tissue samples were cut to 1 mm size and stored in 1 mL of freezing media (10% DMSO (Sigma) and 90% FBS (Gibco)) by slow-freezing in a Nalgene Mr. Frosty freezing container (Sigma). For experimental processing, fresh or cryopreserved samples were made into single cell suspensions by digesting overnight with DNase and collagenase diluted 1:5000 in digestion media (HBSS w/o Ca++ and Mg++, containing 5 mM EDTA and 10mM HEPES) on an orbital shaker. Single-cell suspensions underwent staining for CyTOF as described below. Details for this dataset were described in our published manuscript^23^.

**<u>PBMCs</u>** were isolated from Rhesus macaque blood via a Ficoll gradient. Blood was diluted 1:1 with PBS in a conical tube, and an equal volume of Ficoll was added below the blood layer with a Pasteur pipette. The tube underwent a 30-minute spin with low acceleration and no brake to separate the PBMCs from the rest of the blood contents. The PBMC layer was pipetted out and washed with PBS twice before resuspension in freezing media and slow freezing in a Mr. Frosty for cryopreservation. They were then thawed and underwent staining for CyTOF as described below. Data for this set of samples are publicly available on Cytobank (Beckman Coulter).

### CyTOF experimental pipeline for Human samples

**<u>Placental samples</u>** were collected through the University of Pittsburgh Biospecimen Core as described in Toothaker et al^25^, and from which CyTOF data was extracted. Human placental biopsies were separated by layers for long-term cryogenic storage or immediate experimental processing. They were stored cryogenically and prepared into single-cell suspensions in the same manner as described above for the NHP tissue samples. Single-cell suspensions underwent staining for CyTOF as described below. Details for this dataset were described in our published manuscript^25^.

**<u>PBMCs</u>** were isolated from human blood draws via Ficoll gradient and cryopreserved as described above for the NHP blood samples. Human patients were recruited from Boston Children’s Hospital under the BCH IRB protocol number P00000529. They were then thawed and underwent staining for CyTOF as described below. CyTOF data for this dataset are publicly available on Cytobank (Beckman Coulter).

**<u>CyTOF</u>** staining was performed according to the previously published protocol in Stras et al^26^. Briefly, single-cell suspensions were washed in cell-staining buffer (CSB) composed of PBS with 0.5% bovine serum albumin (Sigma) and 0.02% sodium azide. Viability was assessed with Rh103 (Fluidigm) DNA intercalator. After an additional wash, cells were stained with their respective surface-staining antibody cocktails. For intracellular staining, cells were washed and fixed utilizing FoxP3 fixation and permeabilization kit (Invitrogen). After fixation, cells were washed with the FoxP3 wash buffer and then incubated in their respective intracellular antibody cocktails. Cells were then washed with CSB again and fixed with 1.6% paraformaldehyde (Sigma). After storage in CSB overnight, cells were incubated with 191Ir/193Ir DNA Intercalator (Fludigm) in Maxpar Permeabilization Buffer (Standard Biotools) for cellular identification. On the day of analysis, cells were washed with MilliQ water and resuspended in normalization beads at a 1:10 dilution (Fluidigm). Data collection for the samples was done on a Fluidigm mass cytometer and data were exported as FCS files.

### Data description

To evaluate the gating methods, datasets generated by cytometry technology were collected both from our in-house samples and publicly available datasets. As shown in **Table 1**, six datasets were used in this study: human PBMCs, Rhesus macaque PBMCs, human placental villi^25^, Rhesus macaque placental villi^23^, human bone marrow^27^, and mouse bone marrow^28^. Among these, the first four datasets were generated from our in-house libraries as described above. The human bone marrow^27^ and mouse bone marrow^28^ datasets were collected from the public databases from previous studies. For all six datasets, manual gating and cell population annotation were available (from the original papers for public data, or by our manual gating for in-house data) that can serve as the ground truth for performance evaluation of computer-aided gating methods. Detailed descriptions of each dataset, cell population, and marker table were summarized in **Table 1, Supplementary Table 2 and 3**. Down-sampled datasets to 20,000 cells were generated for tools that could not finish running the full datasets within 3 hours.

**Table 1:** Data Summary.

| Study (Citation) | Sample ID | Manual gating results | Manual gating from 5 experts | Clustering evaluation | Rare population identification | Auto-gating evaluation | Time benchmark evaluation | # Clusters (layer 1, 2, rare) | # Markers (layer 1, 2, rare) | # cells | # study/sample | Species | Tissue |
| --- | --- | --- | --- | --- | --- | --- | --- | --- | --- | --- | --- | --- | --- |
| Human PBMC (CytoBank) | Fresh_Tcell- | Y | Y | Y | Y | Y | Y | 7, 14, 3 | 6, 9, 12 | 190931 | 1 | Human | PBMC |
|  | Fresh_Tcell+ | Y | N | Y | Y | Y | N | 7, 14, 3 | 6, 9, 12 | 171297 | 1 | Human | PBMC |
|  | Frozen_Tcell- | Y | N | Y | Y | Y | N | 7, 14, 3 | 6, 9, 12 | 90382 | 1 | Human | PBMC |
|  | Frozen_Tcell+ | Y | N | Y | Y | Y | N | 7, 14, 3 | 6, 9, 12 | 87459 | 1 | Human | PBMC |
| Rhesus PBMC (CytoBank) | Pooled 6-sample | Y | N | Y | N | Y | N | 6, 13 | 6, 9 | 1252 | 6 | Rhesus | PBMC |
| Human Villi<sup>25</sup> | ID1042 | Y (pooled gating) | N | Y | N | Y | N | 7, 14 | 6, 9 | 8334 | 1 | Human | Villi |
|  | ID1130 | Y (pooled gating) | N | Y | N | Y | N | 7, 14 | 6, 9 | 6485 | 1 | Human | Villi |
|  | Pooled 12-sample | Y | N | Y | N | Y | N | 7, 14 | 6, 9 | 42414 | 12 | Human | Villi |
| Rhesus Villi<sup>23</sup> | ID430 | Y (pooled gating) | N | Y | N | Y | N | 7, 14 | 6, 9 | 70170 | 1 | Rhesus | Villi |
|  | ID437 | Y (pooled gating) | N | Y | N | Y | N | 7, 14 | 6, 9 | 66726 | 1 | Rhesus | Villi |
|  | Pooled 14-sample | Y | N | Y | N | Y | N | 7, 14 | 6, 9 | 345886 | 14 | Rhesus | Villi |
| Human bone marrow<sup>27</sup> | Set1 | Y | N | Y | N | N | N | 8, 20 | 13 | 167039 | 1 | Human | bone marrow |
|  | Set2.H1 | Y | N | Y | N | N | N | 7, 10 | 32 | 72017 | 1 | Human | bone marrow |
|  | Set2.H2 | Y | N | Y | N | N | N | 7, 10 | 32 | 31324 | 1 | Human | bone marrow |
| Mouse bone marrow<sup>28</sup> | S01 | Y | N | Y | N | N | N | 7, 24 | 39 | 53173 | 1 | Mouse | bone marrow |
|  | Pooled 10-sample | Y | N | Y | N | N | N | 7, 24 | 39 | 514386 | 10 | Mouse | bone marrow |

For the human PBMC study, libraries for four treatments were included: fresh unstimulated (-) T cell, fresh stimulated (+) T cell, frozen T cell -, and frozen T cell +. As shown in **Figure 2A** and **Supplementary Table 2 and 3**, the original datasets were based on staining for 52 markers. To serve as ground truth, manual gating was performed at two hierarchical layers. The first layer employed 6 markers (CD3, CD19, CD4, CD8a, CD38, and CD14) to identify 7 major cell populations (CD14- innate, CD38- B cells, Monocytes, NKT cells, CD4 T cells, CD8 T cells, and CD38+ B cells). The second layer further divided the major populations and eventually identified 14 cell types (CD14- HLADR- innate, CD38- B cells, CD4 central memory T cells, CD4 effector memory T cells, CD4 effector T cells, CD4 naïve T cells, CD8 central memory T cells, CD8 effector memory T cells, CD8 effector T cells, CD8 naïve T cells, DCs, Monocytes, NKT, and CD38+ B cells) using 9 markers (CD3, CD19, CD4, CD8a, CD38, and CD14, CCR7, CD45RA, and HLA-DR)^29^. Both layers served as the underlying truth for evaluating computational clustering and auto-gating algorithms. To evaluate the tool’s ability to identify rare populations, innate lymphoid cells (ILCs) and regulatory T cells (Tregs) (each with less than 3% of the total cell numbers) were selected by the manual gating based on 12 markers (CD3, CD19, CD4, CD8a, CD38, and CD14, CCR7, CD45RA, HLA-DR, CD25, FoxP3, and CD127).

**Figure 2:**
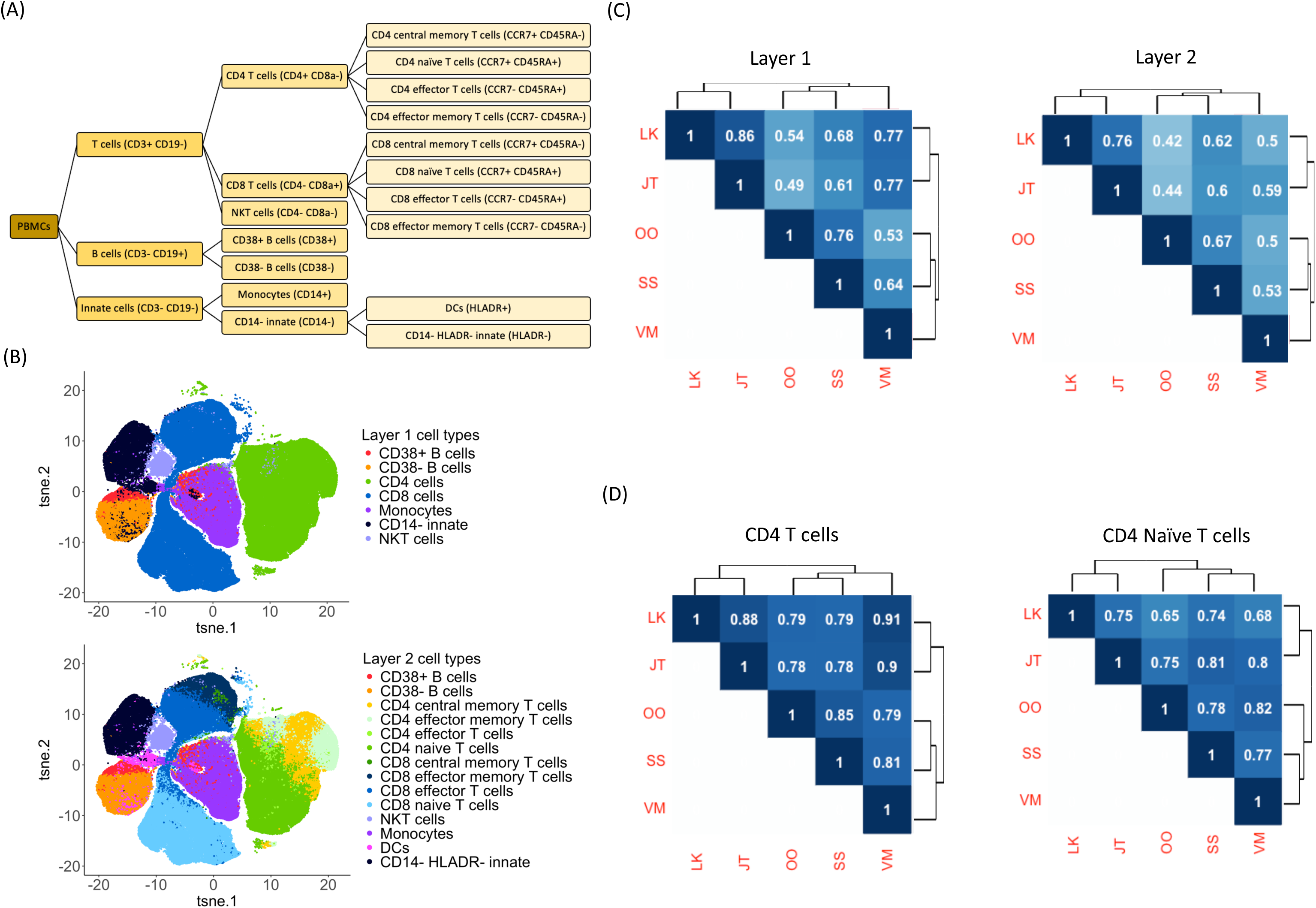
**Coherence of manual gating across five raters**. (A) A hierarchical layer of the human PBMC populations gated by the selected markers. (B) t-SNE figures indicating the manual gating cell population in layer 1 and layer 2 by rater LK. (C) Average pairwise kappa index among five raters with hierarchical clustering to illustrate the similarity of the raters based on their gating assignments. (D) Pairwise kappa index among five raters on CD4 T cells and CD4 Naïve T cells.

In addition to the human PBMCs, the other three in-house datasets were gated into two hierarchical layers. For the NHP PBMC dataset, 6 samples were used and pooled together for the evaluation. Similar to the human PBMC dataset, two hierarchical layers were generated, where the first layer included 6 major populations (CD14- innate, Monocytes, NKT cells, CD8 T cells, CD4 T cells, and CD38+ B cells) using 6 markers (CD19, CD8a, CD14, CD38, CD4, and CD3). The second layer contained 13 clusters (DCs, Monocytes, NKT, CD14- HLADR- innate, CD8 central memory T cells, CD8 effector memory T cells, CD8 effector T cells, CD8 naïve T cells, CD4 central memory T cells, CD4 effector memory T cells, CD4 effector cells, CD4 naïve T cells, and CD38+ B cells) gated by 9 markers (CD19, HLA-DR, CD8a, CD14, CD45RA, CD38, CCR7_CD197, CD4, and CD3). For the human placental villi study, in total 12 samples were analyzed. Two samples (ID1042 and ID1130) with the highest number of cells were selected as representative data. A pooled library merging all 12 samples was also used for evaluation. Similarly, for the NHP villi datasets, a total of 14 samples were analyzed. The top two samples (ID430 and ID437) and the pooled library were compared in this paper. For both human and NHP villi studies, the first layer includes 7 major populations (CD14- innate, Monocytes, NKT cells, CD38- B cells, CD8 T cells, CD4 T cells, and CD38+ B cells) gating by 6 markers (CD18, CD8a, CD38, CD14, CD3, and CD4). The second layer contained 14 cell populations (CD4 central memory T cells, CD4 effector T cells, CD4 effector memory T cells, CD4 naïve T cells, CD8 central memory T cells, CD8 effector T cells, CD8 effector memory T cells, CD8 naïve T cells, DCs, Monocytes, NKT cells, CD38- B cells, CD14- HLADR- innate, and CD38+ B cells) identified by 9 markers (CD19, HLA-DR, CD8a, CD38, CD14, CD45RA, CD3, CCR7, and CD4). **Supplementary Table 2 and 3** describe the cell population and marker table in more detail.

Human and mouse bone marrow datasets were generated from previous studies and downloaded from a public database^27, 28^. For the human bone marrow study^27^, one healthy donor was measured in the first dataset with 13 markers (**Supplementary Table 3**). Manual gating was available with 25 cell populations. To avoid rare populations, several subpopulations with a lower number of cells were merged or removed for our analysis. Eventually, 8 and 20 populations based on manual gating and merging served as the first and second layers of ground truth (**Supplementary Table 2**). In addition, two healthy donor samples were available for the second dataset where 32 markers were measured. On top of the original manual gating, we further removed or merged several smaller populations. Finally, we applied 7 and 10 populations as the first and second layers of underlying truth. For the mouse bone marrow datasets^28^, 10 samples were available in total. We selected two mice (S01 and S02) and a pooled library of 10 samples as representative for the following study. This dataset measured 39 markers and manually gated cells into 7 and 24 populations for the first and second layers. The details of the cellular populations and markers used for gaiting and identification were described in **Table 1** and **Supplementary Tables 2and 3**.

#### Data format, pre-processing, and parameter setting

Cytometry data are commonly stored in a Flow Cytometry Standard (FCS) format with information on both metadata and marker expression. The associated metadata table generally describes experimental and channel information, such as marker name, marker description, and range information. The expression file is in an array or matrix format where each row represents an individual cell, and each column stands for a marker/ channel. These channels correspond to fluorescent markers or heavy metals in flow or mass cytometry data, which have been described in the metadata table^13, 30, 31^.

As the data pre-processing steps, all the negative intensities were trimmed at zero or very small random numbers close to zero (if the algorithms report an error when using multiple zeros as input). Cytometry data were further scaled and inverse-hyperbolic-sine transformed (*X*_*new*_ = asinh(*a* + *b* ∗ *X*_*old*_) + *c*), with a=0, b=0.2 and c=0)^32^. Tools were first applied to the full data with all cells. If the tool could not complete the run within 3 hours, down-sampled data with 20,000 cells were used as an alternative. All the tools were run by default parameter settings except for the number of clusters. If the tool allowed for the specification of the number of clusters to be generated, the true number of clusters was used as input. For detailed scripts for pre-processing and running the tools, please refer to the script files deposited to GitHub: https://github.com/hung-ching-chang/GatingMethod_evalutation/.

### Manual gating collected from five raters

The “fresh T-cells -” library from the human PBMC study was manually gated on the Cytobank^33^ platform by five raters (LK, JT, OO, SS, and VM) who were asked to manually gate the immune cell populations based on their experience independently. No computational algorithm was allowed. Eventually, commonly identified cell populations by all the raters were selected for further evaluation. Based on the hierarchical gating structure in **Figure 2A**, these cell populations were categorized into two layers: the first layer with 7 major populations, and the second layer with 14 subpopulations. The cell populations gated by LK in both layers were visualized by t-Distributed Stochastic Neighbor Embedding (t-SNE)^34^ plots in **Figure 2B**, and the gating results by the other four raters were shown in **Supplementary Figure 1**. To quantitively evaluate the magnitude of agreement between the raters, both kappa index and adjusted Rand index (ARI) measurements were employed, as described in detail in the **Methods** section. A follow-up hierarchical clustering analysis was performed to group the raters with similar gating results based on cellular annotation in the two layers respectively.

### List of unsupervised clustering algorithms evaluated

Unsupervised clustering algorithms group cells into clusters based solely on their marker intensities, lacking the ability to assign the resulting clusters to known cell populations (**Figure 1**). In a previous publication^11^, we summarized 32 tools for unsupervised clustering, which were either specifically designed for cytometry data or for general clustering. Based on the number of input libraries, clustering methods are categorized by the tools that can work on individual samples and tools that need multiple libraries as input. This comparison study only focused on the former ones. As such, we quantitatively compared a total of 22 clustering methods (**Table 2**): ACCENSE^35^, CCAST^36^, ClusterX^30^, Cytometree^37^, densityCUT^38^, DensVM^39^, DEPECHE^40^, FLOCK^41^, flowClust^42^, FlowGrid^43^, flowMeans^44^, flowPeaks^45^, FlowSOM^46^, immunoClust^47^, k-means^48^, PAC- MAN^49^, PhenoGraph^27^, Rclusterpp^50^, SamSPECTRAL^51^, SPADE^52^, SWIFT^53^, and X-shift^28^. These tools have free publicly available software and are compatible with our parameter settings (see **Data format, pre-processing, and parameter setting** section). Manual gating was employed as the underlying truth to evaluate the performance of these unsupervised clustering tools.

**Table 2:** Rank-sum for unsupervised clustering methods.

| Data | Major layer 1 | Major layer 2 | Major layer 1 | Major layer 2 | Rare | Overall ranking | Overall ranking |
| --- | --- | --- | --- | --- | --- | --- | --- |
| Measurement | ARI | ARI | F-measure | F-measure | F-measure | Mean rank | Overall rank |
| ACCENSE | 21 | 21 | 21 | 14 | 13 | 18 | 20 |
| CCAST | 4 | 1 | 8 | 4 | 1 | 3.6 | 2 |
| ClusterX | 18 | 19 | 19 | 20 | 2.5 | 15.7 | 18 |
| Cytometree | 10 | 15 | 6 | 5 | 19 | 11 | 9 |
| densityCUT | 9 | 14 | 18 | 16 | 16.5 | 14.7 | 16 |
| DensVM | 11 | 6 | 9 | 10 | 11 | 9.4 | 7 |
| DEPECHE | 1 | 2 | 5 | 12 | 4 | 4.8 | 5 |
| **FLOCK** | 5 | 5 | 4 | 7 | 7 | 5.6 | 6 |
| **flowClust** | 2 | 8 | 1 | 6 | 6 | 4.6 | 4 |
| **FlowGrid** | 7 | 11 | 13 | 19 | 5 | 11 | 10 |
| **flowMeans** | 15 | 18 | 17 | 15 | 10 | 15 | 17 |
| **flowPeaks** | 12.5 | 16 | 16 | 18 | 8 | 14.1 | 14 |
| **FlowSOM** | 6 | 3 | 2 | 2 | 9 | 4.4 | 3 |
| **immunoClust** | 19 | 20 | 14 | 17 | 20 | 18 | 21 |
| **k-means** | 8 | 9 | 11 | 9 | 18 | 11 | 11 |
| **PAC-MAN** | 3 | 4 | 3 | 1 | 2.5 | 2.7 | 1 |
| **PhenoGraph** | 14 | 12 | 7 | 3 | 12 | 9.6 | 8 |
| **Rclusterpp** | 16 | 7 | 12 | 8 | 16.5 | 11.9 | 12 |
| **SamSPECTRAL** | 17 | 17 | 20 | 21 | 14 | 17.8 | 19 |
| **SPADE** | 20 | 10 | 15 | 13 | 15 | 14.6 | 15 |
| **X-shift** | 12.5 | 13 | 10 | 11 | 21 | 13.5 | 13 |

Note: tool SWIFT can only be applied to human PBMC data, so it's not included in the overall evaluation.

### List of auto-gating algorithms evaluated

Unsupervised clustering methods can identify cell clusters with different characteristics of markers. However, without further annotation of cell populations, these clusters bear no biological insight. Manual annotation of cell clusters is time-consuming and biased. To overcome this problem, several automated annotation and cell-type identification algorithms have been developed. These auto-gating algorithms are designed to identify the resulting clusters from clustering algorithms based on either the prior knowledge of the relationship between lineage markers and the identity of cellular populations or learned from training datasets (**Figure 1**). Our previous publication^11^ summarized six auto-gating algorithms: DeepCyTOF^54^, CyTOF linear classifier^55^, ACDC^56^, MP^57^, OpenCyto^58^, and flowLearn^59^. DeepCyTOF^54^ utilizes deep learning techniques and training data to assign cells to known cell types. Likewise, CyTOF liner classifier^55^ applies training data to predict cell types based on linear discriminant analysis (LDA). In addition, ACDC^56^ uses a prespecified marker matrix to guide the grouping of cells based on a semi-supervised learning approach, and MP^57^ employs a maker matrix to predict cell types through a Bayesian model. In this paper, we compared the performance of the above four methods, while OpenCyto and flowLearn were not evaluated since both of them are not fully automated and require user supervision. As the cell types could be further divided, the underlying truth for cell type identification is controversial and needs to be adjusted depending on the problem at hand. Therefore, to perform a comprehensive evaluation, two hierarchical layers of manually gated references were used in the manuscript.

### Methods for performance evaluation

#### Definition

Cell population identification by manual gating is used as truth to evaluate the performance of unsupervised and supervised algorithms. For each library or pooled library, a set of *n* cells *S* = {*o*_1_, *o*_2_, … , *o*_*n*_} can be grouped into two partitions with *r* populations and *c* populations, defined as *X* = {*X*_1_, *X*_2_, … , *X*_*r*_} and *Y* = {*Y*_1_, *Y*_2_, … , *Y*_*c*_} . For the pairwise comparison between X and Y, the contingency table can be defined as,

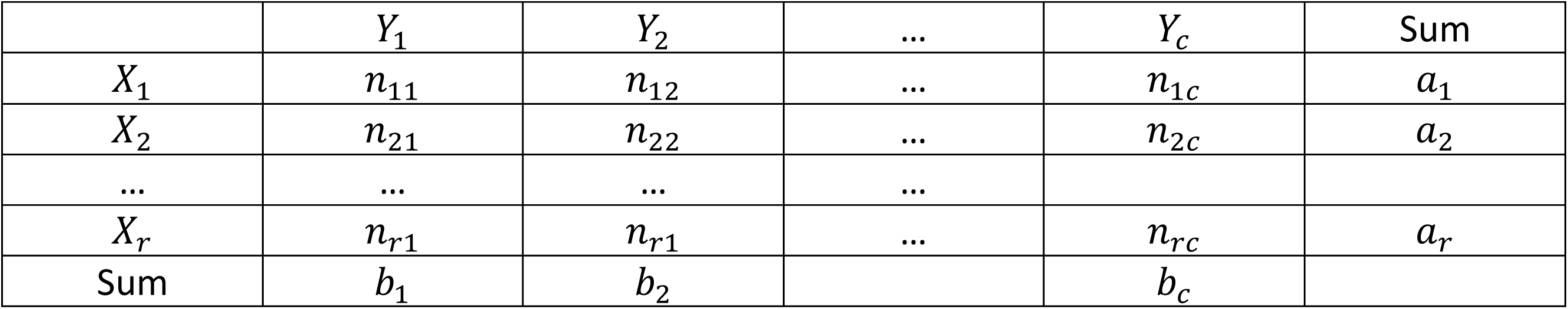

| | $Y_1$ | $Y_2$ | ... | $Y_c$ | Sum |
| --- | --- | --- | --- | --- | --- |
| $X_1$ | $n_{11}$ | $n_{12}$ | ... | $n_{1c}$ | $a_1$ |
| $X_2$ | $n_{21}$ | $n_{22}$ | ... | $n_{2c}$ | $a_2$ |
| ... | ... | ... | ... |  |  |
| $X_r$ | $n_{r1}$ | $n_{r1}$ | ... | $n_{rc}$ | $a_r$ |
| Sum | $b_1$ | $b_2$ | | $b_c$ | |

For *i* ∈ {1, 2, … , *r*} and *j* ∈ {1, 2, … , *c*}, *n*_*ij*_ = |*X*_*i*_ ∩ *Y*_*j*_| represents the number of overlapping cells between partition *X*_*i*_ and *Y*_*j*_, and *a*_*i*_ and *b*_*j*_ indicate |*X*_*i*_| and |*Y*_*j*_|, respectively.

When focusing on a certain population *i* from partition *X* and population *j* from partition *Y*, the 2-by-2 confusion matrix can be written as,

|  |  | Prediction |  |
| --- | --- | --- | --- |
| | | $Y_j$ | $\neg Y_j$ |
| Truth | $X_i$ | TP (true positive) | FN (false negative) |
| | $\neg X_i$ | FP (false positive) | TN (true negative) |

In this paper, we employed the following measurements to indicate the agreement of the two partitions.

**<u>Adjusted Rand Index (ARI)</u>** aims to measure the agreement of two clustering partitions without cell population identification^60^. A value close to 1 indicates high consistency while a value close to zero or even negative means decreased similarity. Assuming *X* to be the true populations by manual gating and *Y* to be the predicted groupings by clustering or auto-gating methods, the ARI is defined as,

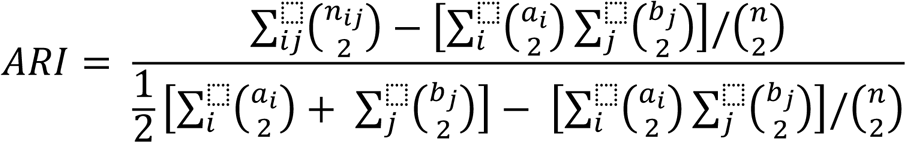

**<u>Kappa index</u>,** also known as Cohen’s kappa coefficient, is an indicator that measures the inter-rater reliability^61^. For binary scenario, the kappa index can be defined as,

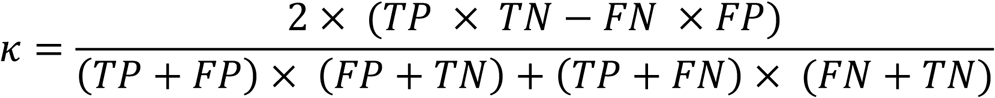

**<u>F-measure</u>** (or F1 score) is an accuracy evaluation measurement to balance the precision and recall in binary conditions^62^. Assuming *X* to be the true population obtained by manual gating and *Y* to be the predicted groupings obtained by clustering or auto-gating methods, precision, recall, and F-measure are defined as follows,

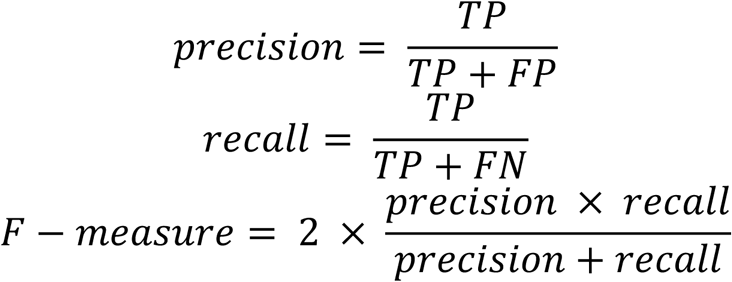

### Computing time evaluation

To benchmark the computing time, the “fresh T cells –” library from the human PBMC dataset was randomly down-sampled to 1000, 2000, 4000, 8000, 16000, 32000, 64000, and 128000 cells. 22 unsupervised clustering and 4 supervised auto-gating methods were applied to this cell number gradient to evaluate both the performance and computing time. For tools where the number of clusters can be set, true number of clusters were set. Otherwise, all the tools were run based on their default parameter settings (supplementary script files submitted to GitHub). Both clustering and auto-gating tools were all run on the same Windows machine (Intel Core i7-8700 CPU @ 3.20GHz 3.19GHz, 16GB RAM, 64-bit operating system, x64- based processor) to reduce machine variability.

## RESULTS

### Manual gating: Consistency across multiple raters

Manual gating has been widely applied to cytometry data to group and annotate individual cells into populations of interest, where researchers have the flexibility to choose the markers and set up the cutoff to define cell populations. However, manual gating is experience-based where researchers tend to gate cells into populations based on their own experiences and marker preferences. Additionally, it’s rather arbitrary when one draws a line to split the populations. To test the consistency of manual gating across different raters, we invited 5 researchers from 3 different labs to perform manual gating on the “Fresh T cells – library” from the human peripheral blood mononuclear cells (PBMC) dataset independently and compared their performances. When excluding the ‘other cells’, cells from the human PBMC study were grouped into 7 major clusters in layer 1 and then further grouped into 14 clusters in layer 2 (**Figure 2A**). **Figure 2B** and **Supplementary Figure 1** illustrated the t-SNE plots of the cell populations gated by rater LK and all the other 4 researchers (JT, OO, SS and VM). To assess the gating similarity among the five raters, the kappa index was used as the evaluation method, with *κ* = 1 indicating complete agreement and *κ* = 0 meaning no agreement. As shown in **Figure 2C**, the pairwise kappa index ranged from 0.44 to 0.86, showing a significant level of variation in manual gating by different experts. The highest kappa index was between LK and JT, 0.86 in layer 1 and 0.76 in layer 2. The lowest kappa index was the one between JT and OO in layer 1 (0.49) and the one between LK and OO in layer 2 (0.42). The kappa indexes in layer 1 were higher than that in layer 2 for the same pair, likely because the extra steps needed for the manual gating for layer 2 introduced the variation. Furthermore, CD4 T cells and CD4 naïve T cells were selected to illustrate the agreement for individual cell populations between the raters in **Figure 2D**. The same patterns were observed, though the actual numbers varied.

Given the raters were only instructed to gate the cell population by their own experience, the raters not only gated common populations differently but also focused on different sets of subpopulations. For example, LK gated 39 subpopulations in total, while OO gated 24 subpopulations, where FoxP3^+^ CD4 T-cells and GATA-3^+^ CD4 T cells were gated only by LK, while Th1 CD4 T cells and Th17 CD4 T cells were gated only by OO (**Supplementary Table 4**). We observed that the lines drawn to split the populations were subjective, and the raters had their own preferences in splitting the populations. These results gave rise to the need for more reproducible and less labor-intensive gating strategies, which motivated us to review the clustering and auto-gating tools in the following sections.

### Clustering: Detection of major cell populations

Unsupervised clustering refers to the computational method for cell grouping without population annotation. 32 unsupervised clustering tools were reviewed and discussed in our previous study^11^. When we attempted to evaluate them comprehensively, 22 tools could be applied to individual datasets and installed successfully on our end (**Table 2**). To evaluate the performance of unsupervised clustering algorithms, we applied them to six datasets as described in **Table 1**. For each study, the major populations were identified hierarchically in two layers: layer 1 represented top-level major populations and layer 2 included lower-level and detailed populations. Both adjusted Rand index (ARI) and F-measure (described in the **Methods** section) were applied to evaluate the agreement between the clustering results for each tool and the manual gating truth. The ARI values for each dataset and method were illustrated in heatmaps with rows representing the tools and columns indicating datasets (**Figure 3 A-B**), where ARI = 1 indicated an exact match between the two groups, while an ARI value close to zero meant a lack of consistency. As illustrated by the heatmap color for ARI, results for both layers 1 and 2 indicated overall high performance for the human PBMC and human bone marrow and mouse bone marrow studies, while comparatively lower performance for the rhesus PBMC, human placental villi, and rhesus placental villi were observed. This was potentially due to human antibodies suboptimal staining of rhesus samples or the need for altered phenotyping in rhesus samples. Additionally, staining within the tissue (placental villi) as compared to blood resulted in less distinct marker differences than that obtained from PBMCs.

**Figure 3:**
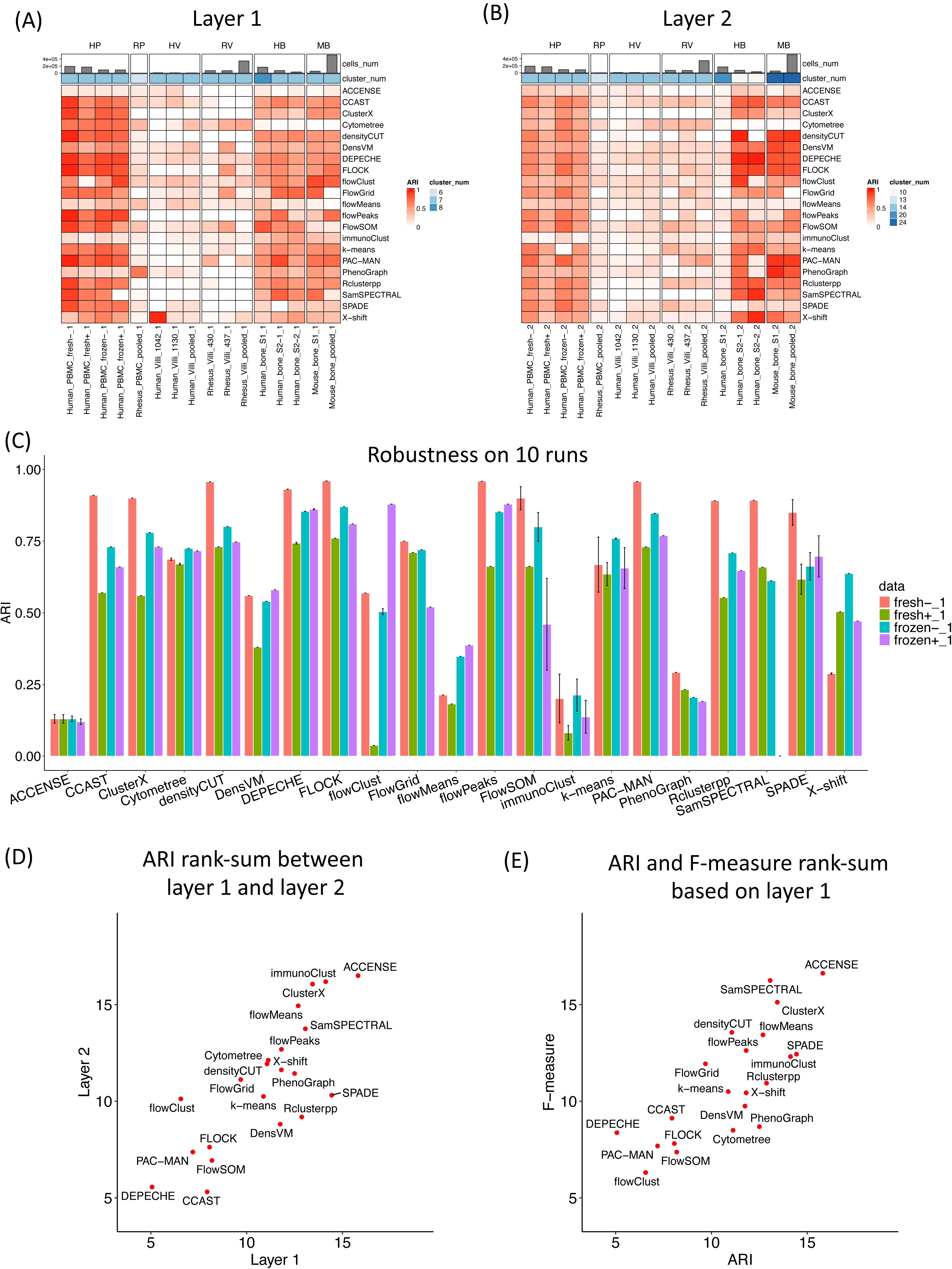
Major population detection by unsupervised clustering algorithms. (A) Performance of the clustering algorithms based on layer 1 cell populations using the ARI measure. (B) Performance of the clustering algorithms based on layer 2 cell populations using the ARI measure. (C) Robustness of the clustering algorithms on 10 runs using the ARI measure. (D) Rank-sum between layer 1 and layer 2 using the ARI measure. (E) Rank-sum between the ARI and F-measure based on layer 1 data.

Many of the clustering algorithms include random initial steps to learn the grouping and subsequently cluster cells, which will result in different clustering outcomes for different runs. To check the robustness/ consistency across multiple runs, all the tools were applied to the human PBMC dataset for 10 independent runs. As shown in **Figure 3C**, the bar plot indicated the average and standard deviation of the 10 repeated runs. Among these clustering algorithms, tools such as ACCENSE, FlowSOM, immunoClust, k-means, and SPADE presented larger variations among multiple runs, while the remaining tools generated comparatively consistent or even identical results.

When comparing the tool’s performance in detecting top-layer major populations and lower-layer detailed populations, we evaluated their performance against two hierarchical truths (described in the **Method** section). For each truth layer, the rank-sum of the tool over multiple studies was calculated and averaged (**Supplementary Table 5**). Eventually, the average rank-sum per layer was compared (**Figure 3D)** and the overall ranking was summarized in **Table 2**, where lower rank indicated higher consistency with the truth. When comparing tool performance between layer 1 and layer 2, **Figure 3D** showed high agreement for most of the tools (close to the diagonal line), except for DensVM, Rclusterpp, and SPADE. These three tools presented comparatively lower performance in layer 1 but higher performance in layer 2 (higher rank in layer 1 than layer 2). These tools tended to provide a larger number of clusters in their default parameter settings, which resulted in better detection of more detailed sub-cell populations than the major ones.

In addition to the ARI measure, the F-measure was also applied in our study as an alternative evaluation benchmark (see **Methods** section). Per tool-predicted cluster, the highest F-measure was applied as the best match between the predicted cluster and the true cluster. Eventually, averaged F-measures were calculated to indicate the overall performance of the clustering algorithms. **Supplementary Figure 2** illustrated tool performance quantified by F-measure and **Table 2** summarized their overall performance ranking. When comparing the performance consistency between multiple measurements, **Figure 3E** showed the average rank-sum of the ARI and F-measure. The majority of the tools showed high agreement between the two measurements, while some tools showed comparatively higher performance for one measurement than the other. For example, PhenoGraph, DensVM, Rclusterpp, and SPADE resulted in better performance by the F-measure but lower performance by the ARI measure. This may be due to the mapping between tool-predicted clusters with the ground truth. In general, a tool yielding a larger number of clusters than the truth will tend to get a better performance by F-measure, which will choose the best cluster to match with the truth. However, this tool will receive lower performance by ARI for punishing a large number of clusters. In conclusion, when considering both layer and measurement effects, DEPECHE, CCAST, FlowSOM, PAC-MAN, and FLOCK performed overall the best for the selected dataset (**Table 2** and **Supplementary Table 5**).

### Clustering: Computing time benchmarking

Besides clustering accuracy, computational cost is another factor in evaluating these tools. In this study, we selected the “human PBMC fresh T cells - library” as an example for benchmarking the computing time. Specifically, we subsampled cells from this dataset by log2 gradient numbers: 1k, 2k, 4k, 8k, 16k, 32k, 64k, and 128k. All the clustering algorithms were applied to these eight subsets based on default parameter settings to record their computing times. **Figure 4A** demonstrated the computing time (average of 10 runs) per tool per data subset. Among these tools, k-means, FlowGrid, and densityCUT overall consumed a shorter time for these datasets. To predict the running time for a larger number of cells, we fitted the computing time to a logistic regression in terms of the number of cells (**Figure 4B** and **Supplementary Figure 3**). Except for flowPeaks (**Figure 4B**) which consumed constant computing time across different datasets, all the other tools (for example, DensVM in **Figure 4B**) resulted in longer computing time with an increasing number of cells. However, these tools yielded different increases in computing time for additional cells, which were reflected by the slope of the regression model. Tools such as FLOCK, FlowSOM, and FlowGrid achieved comparatively shorter computing costs for additional cells (with slope to be 0.12, 0.25, and 0.43 in **Supplementary Figure 3**), indicating better compatibility to analyze larger datasets. In contrast, tools such as ClusterX and CCAST increased the amount of computing time fast (with slope to be 1.89 and 1.52 in **Supplementary Figure 3**), which may result in much longer computing time for larger libraries.

**Figure 4:**
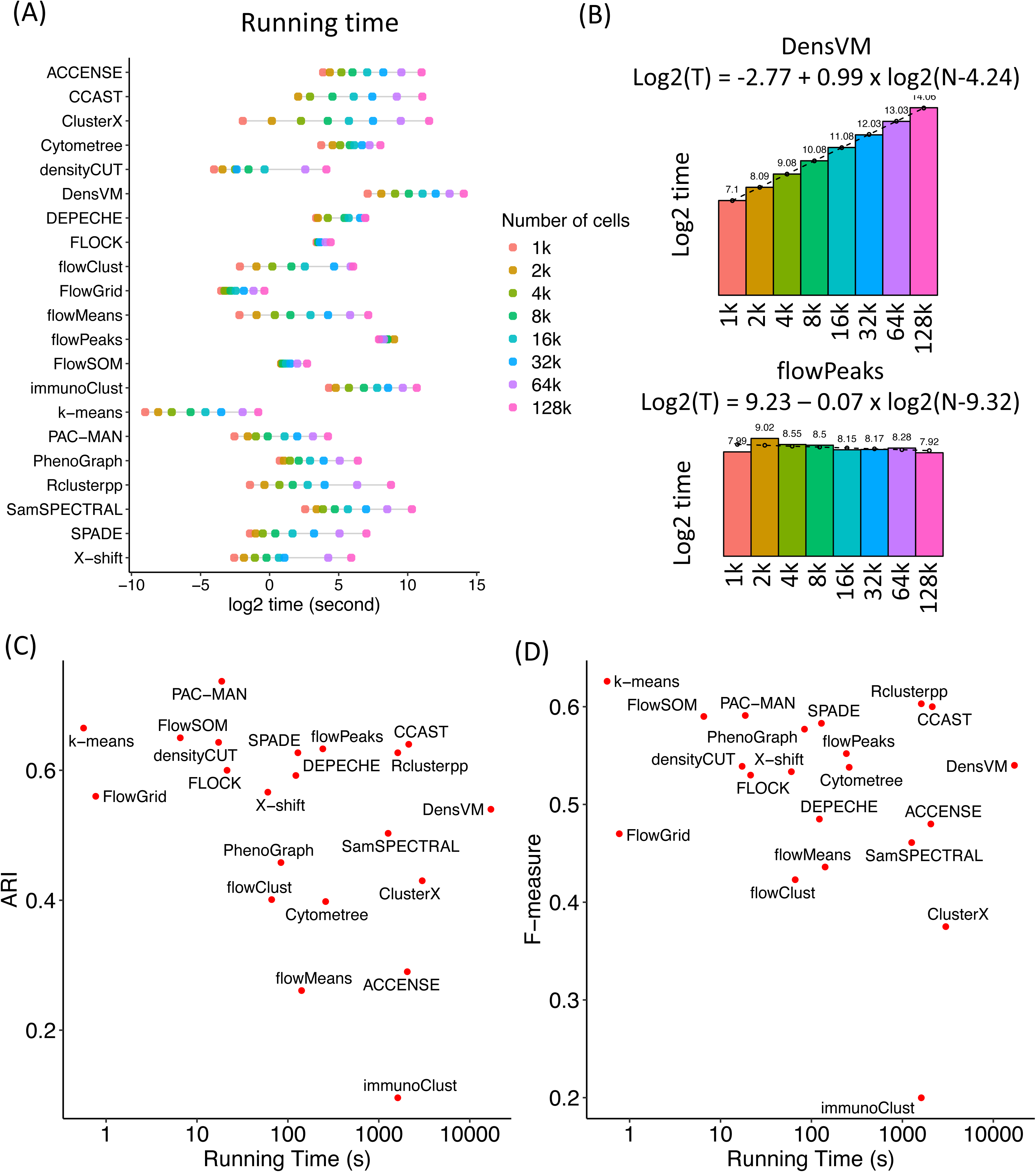
Time benchmark for unsupervised clustering algorithms. (A) Time benchmark for clustering algorithms on gradient number of cells. (B) Computing time for two representative algorithms with linear (DensVM) and flat (flowPeaks) increasing speed. Bar-graph represents the real running time and the dashed line for the predicted running time when fitting into the regression model. (C) The comparison between computing time and ARI performance. (D) The comparison between computing time and F-measure performance.

For an overall evaluation of the tools, **Figures 4C** and **4D** visualized the tool performance and the computing time simultaneously. In **Figure 4C**, tools PAC-MAN, k-means, FlowSOM, densityCUT, FLOCK, and FlowGrid have high ARIs with comparatively shorter computing time, while in **Figure 4D**, tools k-means, FlowSOM, PAC-MAN, PhenoGraph, densityCUT, X-shift, and FLOCK yielded the best performance quantified by F-measure and lower computing cost.

### Clustering: Detection of rare populations

We referred to the ‘rare population’ as the cell type with a lower number of cells compared with the major populations. Since each rare population only represents less than 3% of the whole library, it is easily missed when all the cells are clustered. To evaluate the capability of the clustering algorithms to identify rare populations, we selected two rare populations from the human PBMC dataset to illustrate (**Figure 5A**): innate lymphoid cells (ILCs) and regulatory T cells (Tregs). The number of cells within each population and the markers applied were listed in **Supplementary Table 2 and 3.** Their distributions to the whole libraries were shown in the t-SNE plot in **Figure 5A**. When aiming for rare population detection, parameters of the clustering tools were adjusted to detect a larger number of small clusters, rather than big clusters for the major population. The detailed settings were shown in the supplementary script files. F-measure was applied to evaluate the consistency between the clustering algorithms and the manual gating truth. As shown in **Figure 5B**, the heatmap visualized the performance of 22 clustering algorithms in detecting these two rare populations across the four human PBMC libraries. In general, many tools resulted in overall high and robust performance across multiple libraries and for both of the rare populations, including ACCENSE, CCAST, ClusterX, DensVM, DEPECHE, FLOCK, flowClust, FlowGrid, flowMeans, flowPeaks, FlowSOM, PAC-MAN, and PhenoGraph.

**Figure 5:**
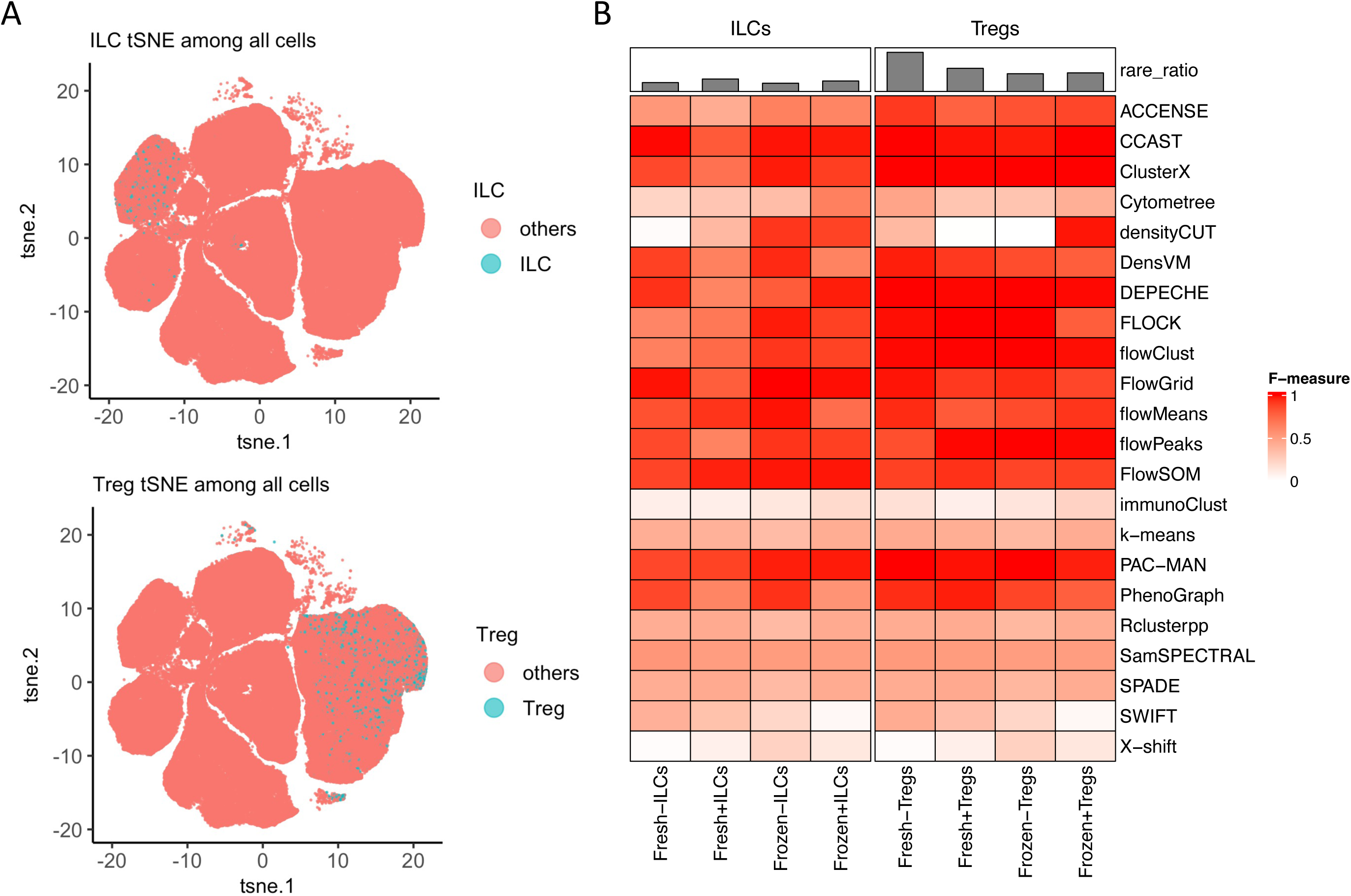
Rare population detection by clustering algorithms. (A) Manual gating for the selection of the two rare populations: ILCs and Tregs. (B) Heatmap for the performance of the clustering algorithms on the rare population based on the F-measure.

### Auto-gating: Prior knowledge preparation and performance evaluation

Supervised or semi-supervised auto-gating methods refer to the algorithms that take in prior knowledge or training sets to train the model, and then perform cell population identification based on these parameters. In this paper, we quantitatively reviewed four auto-gating methods: DeepCyTOF^54^, CyTOF linear classifier^55^, ACDC^56^, and MP^57^. To check the agreement between these auto-gating algorithms with the manual gating truth, both ARI and F-measure were applied to evaluate the performance of these tools on the PBMC and the placental villi data. As shown in **Figure 6A**, the heatmap indicated the ARI value for each tool (row) across each dataset (column) when using the layer 1 manual gating as truth. Across all the datasets, DeepCyTOF and CyTOF Linear Classifier had an overall better performance, while ACDC and MP only performed well with the human PBMC data. The potential reason might be that the Human PBMC dataset had a higher number of cells to cover a more robust set of different immune cell populations when compared with the other datasets. In addition to layer 1, layer 2 truth was applied and demonstrated similar patterns (**Figure 6B)**. The rank-sum comparison between layers 1 and 2 was shown in **Figure 6C**, where these four tools showed a high agreement of performance when applied to the two hierarchical truths, and DeepCyTOF and CyTOF Linear Classifier achieved high ARI consistently. To evaluate the robustness of these tools, we ran all the algorithms 10 times on the Human PBMC data. As shown in **Figure 6D**, all four auto-gating tools presented low variations across multiple runs. Besides the ARI measurement, a similar performance evaluation was measured by the F score and the same conclusion can be drawn (**Supplementary Figure 4)**. **Table 3** summarized the rank-sum per measurement and their overall ranking, where DeepCyTOF and CyTOF Linear Classifier were the top two algorithms recommended based on our evaluation.

**Figure 6:**
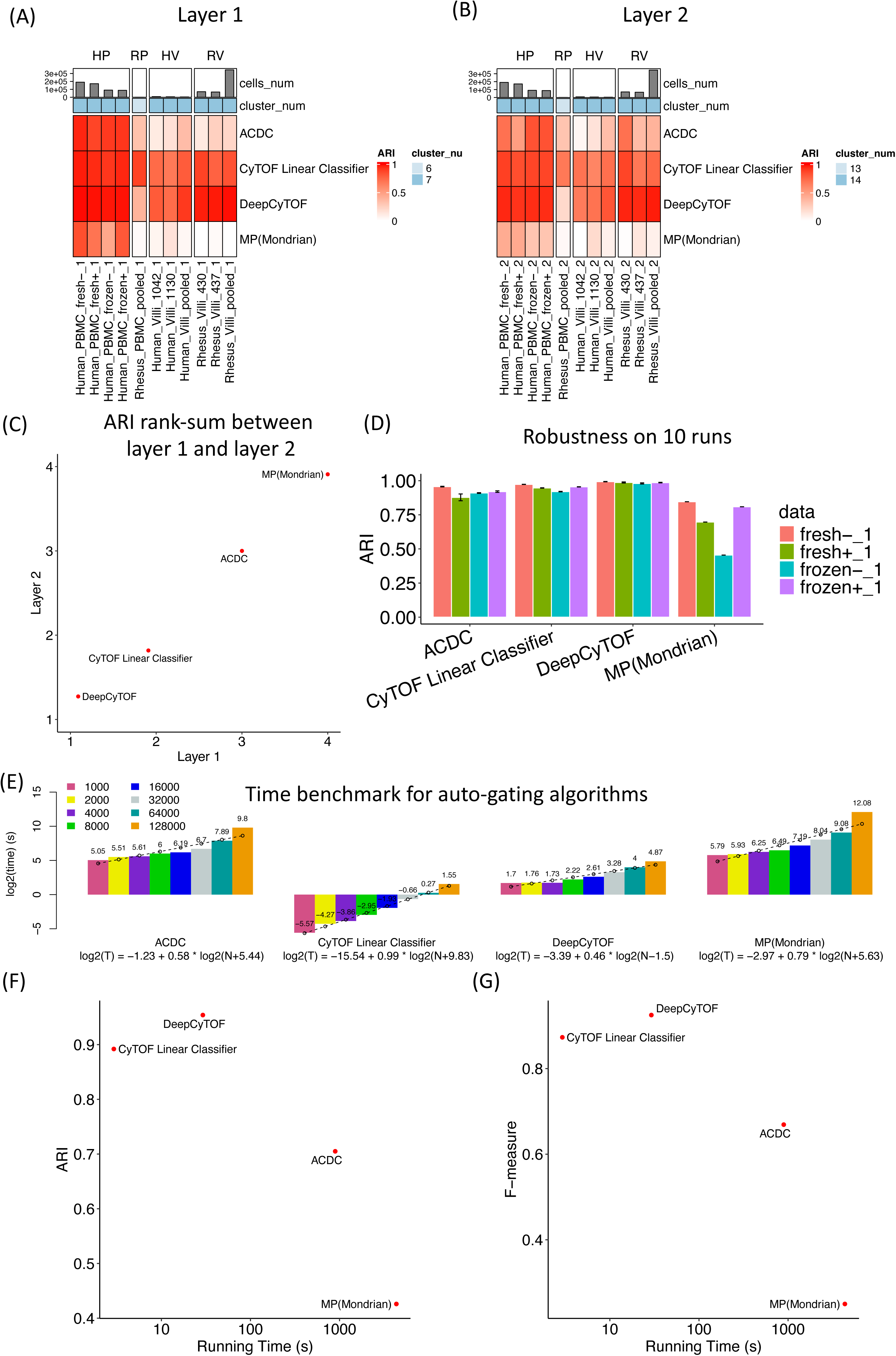
Population detection by supervised auto-gating algorithms. (A) Performance of the auto-gating algorithms based on layer 1 cell populations using the ARI measure. (B) Performance of the auto-gating algorithms based on layer 2 cell populations using the ARI measure. (C) Rank- sum between layer 1 and layer 2 using the ARI measure. (D) Robustness of the auto-gating algorithms on 10 runs using the ARI measure. (E) Time benchmark for auto-gating algorithms on gradient number of cells. Bar-graph represents the real computing time and the dashed line for the predicted computing time when fitting into the regression model. (F) The comparison between computing time and ARI performance. (G) The comparison between computing time and F-measure performance.

**Table 3:** Rank-sum for supervised clustering methods.

| **Data** | **Major layer 1** | **Major layer 2** | **Major layer 1** | **Major layer 2** | **Overall ranking** | **Overall ranking** |
| --- | --- | --- | --- | --- | --- | --- |
| **Measurement** | **ARI** | **ARI** | **F-measure** | **F-measure** | **Mean rank** | **Overall rank** |
| **ACDC** | 3 | 3 | 3 | 3 | 3 | 3 |
| **CytoF Linear Classifier** | 2 | 2 | 1 | 1 | 1.5 | 1.5 |
| **DeepCyTOF** | 1 | 1 | 2 | 2 | 1.5 | 1.5 |
| **MP (Mondrian)** | 4 | 4 | 4 | 4 | 4 | 4 |

In the next step, we aimed to check the computing cost of these auto-gating tools. **Figure 6E** indicated the computing time across a gradient number of cells for each of the tools used (similar pipeline as was done for the clustering methods). DeepCYTOF achieved a low computing time and the lowest increasing slope (0.46) for the regression model among the four tools. CyTOF Linear Classifier resulted in the overall shortest computing time while maintaining within linear slope (0.99). When considering the tool’s computing time and performance at the same time, **Figures 6F** and **6G** illustrated that both DeepCyTOF and CyTOF Linear Classifier achieved overall higher performance and shorter computing cost.

## DISCUSSION

This paper comprehensively investigated and compared three main categories of methods for analyzing cytometry data: manual gating, unsupervised clustering, and supervised auto-gating (**Figure 1**). Among them, manual gating involves visually inspecting multidimensional plots of the data and drawing boundaries (gates) around populations of interest. This gating method is widely applied by expert researchers and can be operated with flexibility and transparency. However, manual gating has limitations when dealing with high-dimensional and large-scale datasets, and the results are subject to the researcher’s experience. In contrast to manual gating, unsupervised clustering and auto-gating methods are computer-assisted algorithms, which have the advantages of automation and reproducibility for large datasets, but face the limitations in transparency and parameter sensitivity. When distinguishing these two computational gating methods, clustering algorithms aim to group similar cells into clusters without predefined populations, while auto-gating tools are designed to mimic the manual gating process to identify and gate cell populations. Although many automated gating algorithms have been developed, manual gating and unsupervised clustering followed by manual annotation are the most widely used pipelines for cell population identification.

In this manuscript, we systematically evaluated 22 unsupervised clustering algorithms and 4 auto-gating tools (supervised or semi-supervised). **Table 2 and 3** summarized the rank of these tools when applied to six cytometry datasets by both ARI and F-measure benchmarks and based on both layers 1 and 2 truth. **Figure 7** presented a workflow and comparison for an overall recommendation of the algorithms. Among all the computer-assisted tools, if no prior knowledge nor training data was available, unsupervised clustering tools were suggested. Among them, tools with higher performance and shorter computing time were recommended. Other specific recommendations were made as well. For example, if the users wanted to specify the number of clusters, tools such as PAC-MAN, FlowSOM, flowClust, DEPECHE, k-means, X-shift, flowPeaks, and SPADE would be good options. If the researchers were interested in graphical visualization of the clustering results, tools such as FlowSOM, Cytometree, SPADE, X-shift, and PhenoGraph could yield figures for hierarchical trees or gating networks. If the rare populations were the major focus of the study, we would recommend DEPECHE, FLOCK, FlowGrid, flowPeaks, FlowSOM, PAC- MAN, and PhenoGraph. When considering all these factors and balancing accuracy and computing time, PAC-MAN and FlowSOM were the top two tools to be recommended. In addition to clustering algorithms, auto-gating methods were suggested for studies with prior knowledge of cell populations. In general, our evaluation study suggested DeepCyTOF and CyTOF Linear Classifier as they had an overall higher accuracy and shorter computing time.

**Figure 7:**
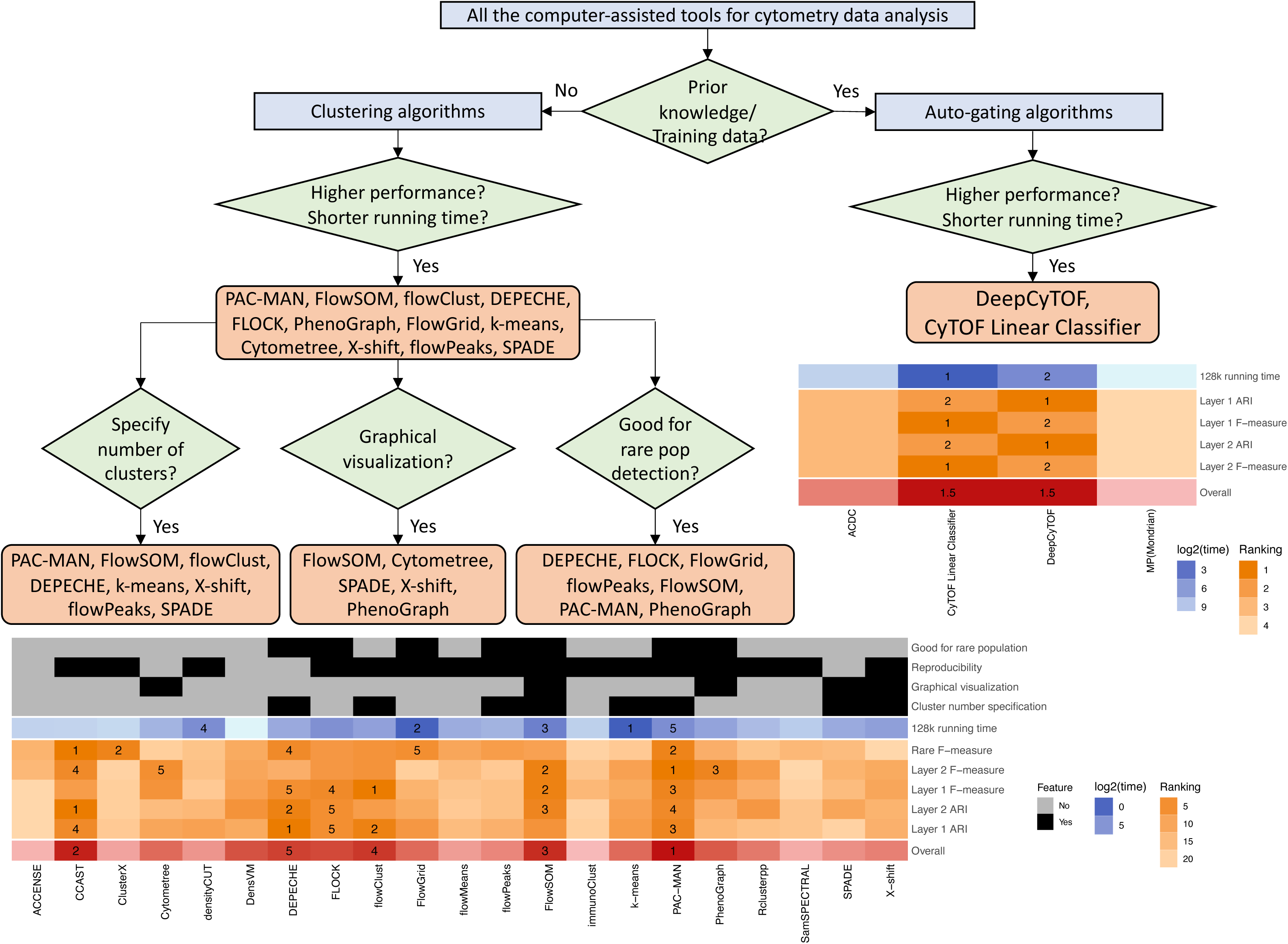
Overall recommendation for the selection of computer-assisted algorithms. Workflow: recommendation of the cytometry tools for different applications. Heatmap for clustering tools: including tool features (black), rankings for computing time (blue), accuracy evaluation (orange), and overall accuracy ranking (red). Heatmap for auto-gating tools: rankings for computing time (blue), accuracy evaluation (orange), and overall accuracy ranking (red).

Depending on different research interests, some studies only focus on major cell types, while other studies aim to explore detailed subpopulations or even rare populations. In this study, we provided two hierarchical layers of truth: layer 1 for major populations and layer 2 for detailed types. When comparing the consistency of the gating results between the two layers, most of the tools had high agreement (**Figures 3D** and **6C**), while tools that generally yield a higher number of clusters tended to favor layer 2 to detect more detailed populations. As an overall suggestion, if the researchers are interested in subpopulations or even rare populations, tools that can specify the number of clusters or perform well for rare populations are recommended (**Figure 7**).

Among the six datasets, diverse samples were employed for tool comparison, including multiple tissue types (PBMCs, bone marrows, and placental villi), fresh and frozen samples, and samples from different species (human, non-human primate, and mouse). In general, PBMC and bone marrow samples outperformed the placental villi. This is likely due to marker expression on cells within the tissue being less distinct than in the bone marrow and the peripheral blood. As such, auto-gating for tissue samples is not recommended. In contrast, the difference between fresh and frozen PBMC samples was trivial, which indicated that this treatment wouldn’t influence the antibody capture in the cytometry experiment^24^. In addition, samples from three species were included in this study: human, rhesus, and mouse. The human samples performed the best, likely secondary to more optimized antibodies and better phenotyping.

While this study aimed to provide a comprehensive evaluation of the cytometry gating methods, several limitations still needed to be addressed. First, given that clustering and auto-gating tools update quickly, we’ve tried our best to apply the newest version of the software. Tools that were no longer maintained, failed to install on our computer, or could not complete running on some datasets were not included in the final results. Second, although we attempted to include diverse datasets, the comparison of the results may differ by the specific datasets applied. In this study, both our in-house data and public datasets were employed to provide an unbiased evaluation. Third, the truth was generated by the manual gating of the most experienced research in our evaluation. As such, we only focused on the well-defined immune cell populations for the purpose of this study. Lastly, the comparison of auto-gating methods on rare populations was not performed, given the limitation of prior knowledge and training data.

## DATA AVAILABILITY

All animals analyzed in this study are stored in accordance with Institutional Animal Care and Use Committee (IACUC) guidelines at the University of California Davis and endorsed by the University of California, Los Angeles (protocol #20330 and #22121). Requests can be directed to Liza Konnikova. Human blood samples were collected at Boston Children’s Hospital under Institutional Review Board (IRB) protocol IRB-P00000529, and placental villi samples were collected under IRB approval at the University of Pittsburgh. Cytometry data for our in-house datasets are publicly available on Cytobank (Beckman Coulter). All the supplementary script files are available at GitHub: https://github.com/hung-ching-chang/GatingMethod_evalutation.

## ACKNOWLEDGEMENT

We thank Stephanie Stras for helping with manual gating.

## AUTHOR CONTRIBUTIONS

JT, LK, EGS, BM, and VM performed the cytometry experiments. LK, JT, OO, Stephanie S, and VM performed the manual gating on the in-house datasets. PL and LK computationally analyzed the manual gating data. SL mined the public cytometry datasets. PL, YF, XX, JZ, and SL performed the clustering algorithms, and PL and YP explored the auto-gating pipelines. SBS oversaw the human PBMC dataset, and PP and SK oversaw the NHP. PL, YP, HC, JL, and SL integrated all the results and generated the figures and tables. PL, GT, LK, and SL wrote the manuscript. SL, LK, and GT oversaw the whole study.

## SUPPLEMENTARY FIGURE LEGEND

**Supplementary Figure 1:**
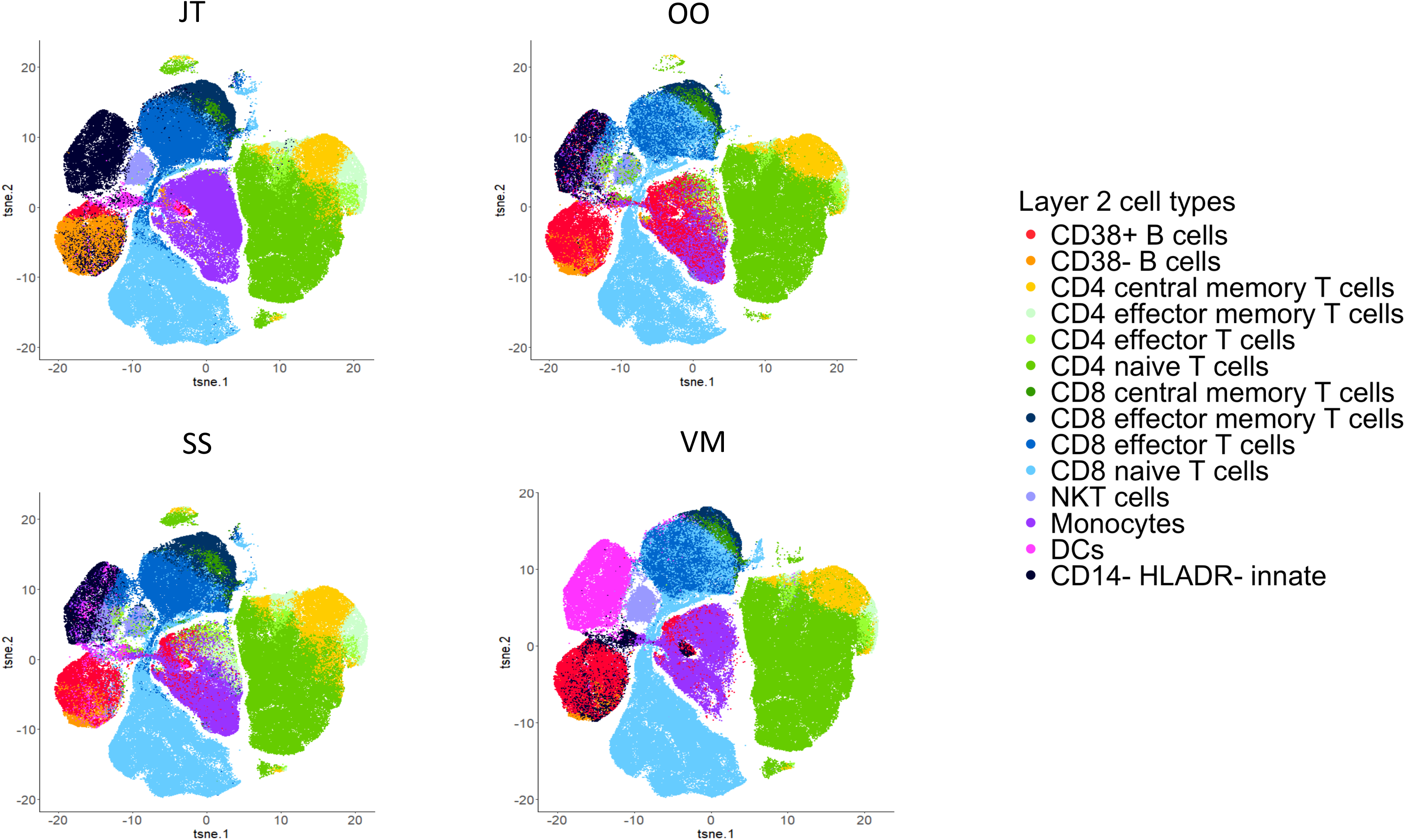
t-SNE figures indicating the manual gating cell population in layer 1 and layer 2 by rater JT, OO, SS, and VM.

**Supplementary Figure 2:**
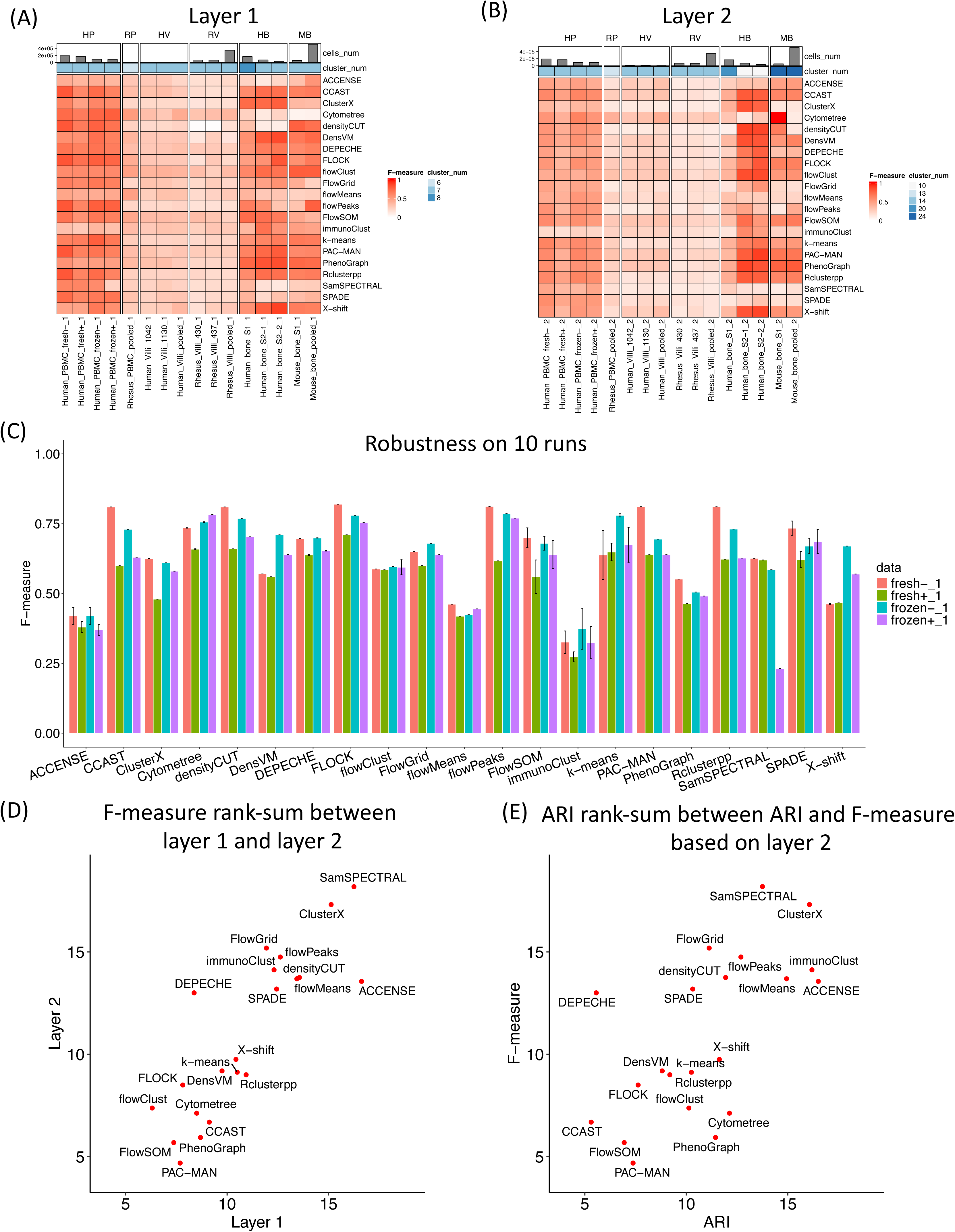
The detection of major populations by unsupervised clustering algorithms, quantified using the F-measure. (A) Performance of the clustering algorithms based on layer 1 cell populations using the F-measure. (B) Performance of the clustering algorithms based on layer 2 cell populations using the F-measure. (C) Robustness of the clustering algorithms on 10 runs using the F-measure. (D) Rank-sum between layer 1 and layer 2 using the F-measure. (E) Rank-sum between the ARI and F-measure based on layer 2 data.

**Supplementary Figure 3:**
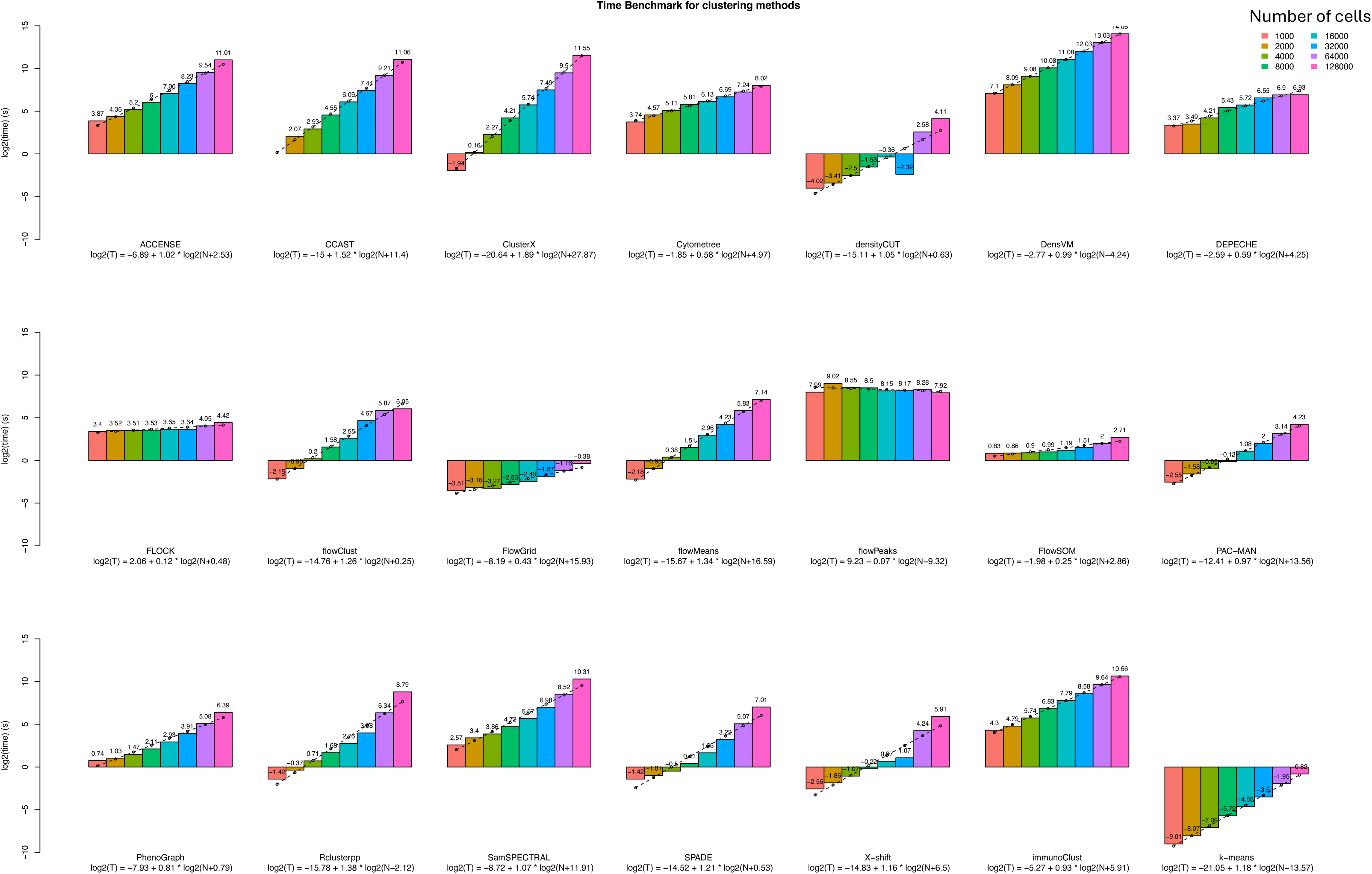
Computing time for the clustering algorithms on a gradient number of cells. Bar-graph represents the real computing time and the dashed line for the predicted computing time when fitting into the regression model.

**Supplementary Figure 4:**
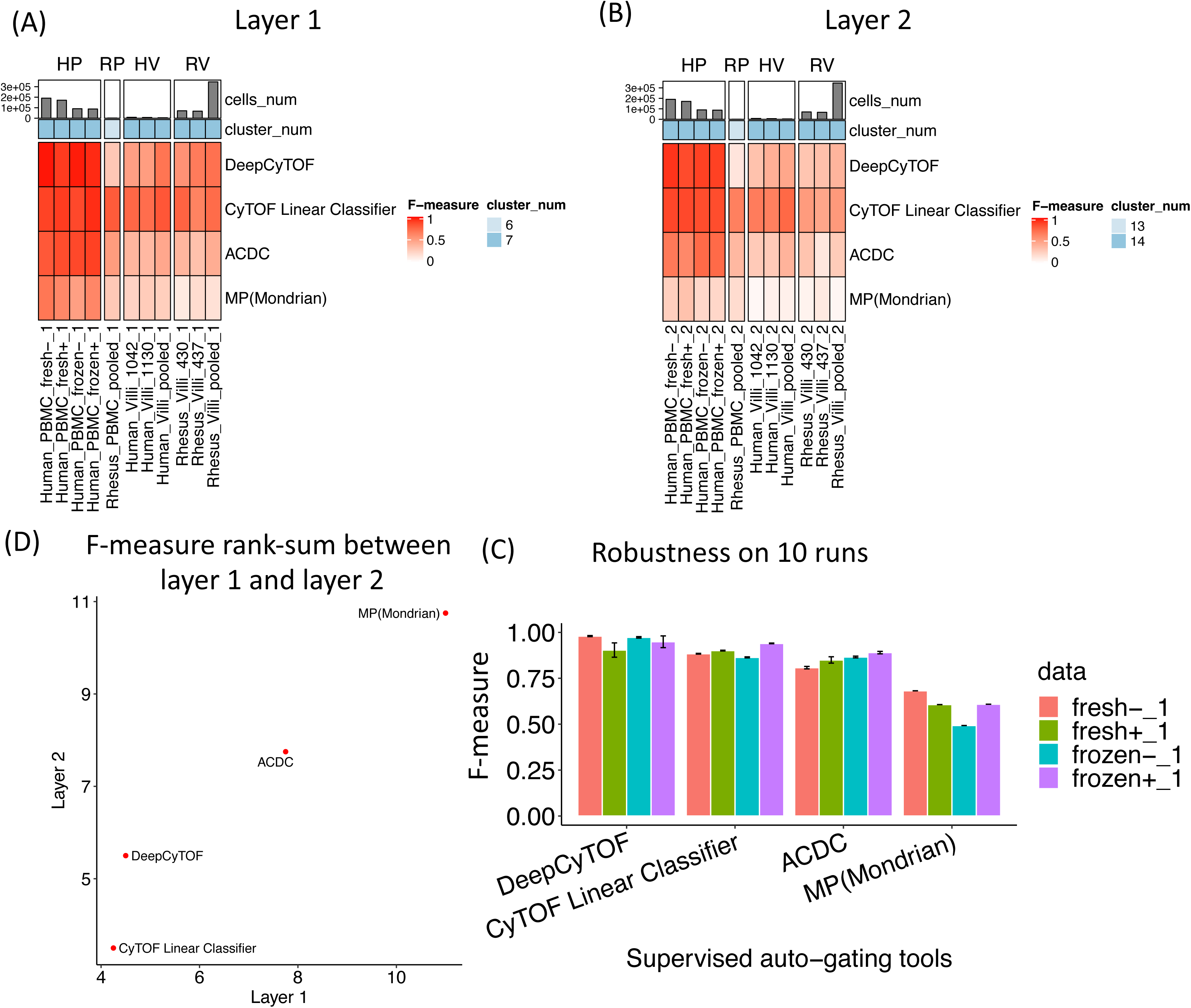
Population detection by supervised auto-gating algorithms quantified using F-measure. (A) Performance of the auto-gating algorithms based on layer 1 cell populations using the F-measure. (B) Performance of the auto-gating algorithms based on layer 2 cell populations using the F-measure. (C) Rank-sum between layer 1 and layer 2 using the F-measure. (D) Robustness of the auto-gating algorithms on 10 runs using the F-measure.

## SUPPLEMENTARY TABLE

Supplementary Table 1: Comparison between the published review paper and our study.

Supplementary Table 2: Number of cells per cell population.

Supplementary Table 3: Marker table.

Supplementary Table 4: Manual gating details by the five raters.

Supplementary Table 5: Rank and rank-sum for the unsupervised clustering methods.

Supplementary Table 6: Rank and rank-sum for the supervised auto-gating methods.

